# Ocean acidification changes diet effects and differentially impacts two populations of red abalone (*Haliotis rufescens*)

**DOI:** 10.64898/2026.06.18.733263

**Authors:** Sara E. Boles, Daniel S. Swezey, Kristin M. Aquilino, Haley K. Stott, Laura Rogers-Bennett, Doug Bush, Eric Sanford, Andrew Whitehead

## Abstract

Absorption of CO_2_ by global oceans is decreasing pH resulting in ocean acidification (OA). Impacts on shellfish have been documented in ecologically and commercially important species. We examined the influence of diet and OA between two populations of red abalone (*Haliotis rufescens*) a species of aquaculture importance and declining wild populations. Populations experience different exposure histories: strong upwelling (Van Damme, California [VD]) historically exposed to low-pH conditions and weak-intermittent upwelling (Santa Barbara, California [SB]). Abalone were cultured under control-pH or OA-conditions and fed crustose coralline algae (CCA) or diatoms used in aquaculture. We tested treatment effects of population, settlement diet, and OA-exposure on survival as influenced by larval-energy stores. Survival in both populations was enhanced by CCA when cultured under both treatment conditions; however, by later stages, this effect remained only for SB. SB had reduced post-settlement survival when cultured under OA-conditions, whereas post-settlement survival of VD was not. Diet affected the relationship between larval-energy and post-settlement survival; a positive relationship when fed diatoms and a negative relationship with CCA. The relationship between larval energy and post-settlement survival was stronger in VD. CCA enhanced juvenile growth in SB cultured abalone at both three-months and one-year post-settlement. Settlement diets can reduce the impacts of OA on early-life stages of abalone, but population differences driven by underlying energetics affect the consistency of this outcome. These findings illuminate the impacts from OA, suggesting populations may be at risk, and inform strategies for developing and sustaining shellfish aquaculture in the face of changing ocean conditions.

## Introduction

Anthropogenically-emitted carbon dioxide (CO_2_) absorbed by the world’s oceans has resulted in a decrease in global ocean pH, a process known as ocean acidification (Caldeira & Wickett, 2003). Accelerating CO_2_ emissions are projected to cause a decrease in oceanic pH by 0.3 to 0.4 pH units, increasing the acidity [H^+^] by 100 to 150% by the year 2100 (Caldeira & Wickett, 2003; Orr et al., 2005). OA impacts on shellfish include decreased growth (Watson et al., 2009; Barton et al., 2012), reproduction (Baros et al., 2013; Boch et al., 2017), survival (Talmage & Gobler, 2010; Crim et al., 2011), and nutritional health (Tate et al., 2017; Lemasson et al., 2019). These consequences threaten the persistence of wild populations and the sustainability of shellfish aquaculture production. Further, OA threatens the value of commercial fisheries and the quality of seafood in the broader global commercial shellfish supply chain (Gazeau et al., 2007) valued at >$281.5 billion USD in 2020 (FAO, 2022).

Among mass produced shellfish species, abalone (*Haliotis* spp.) represents one of the most desired and expensive luxury seafood items currently on the market, with an annual global production value estimated at $3 billion USD in 2016 (Cook, 2016). The harvest of abalone by humans spans a long and complex history, and abalone hold significant food and cultural value for many indigenous peoples around the world (Glassow, 1987; Vellanoweth et al., 2006; Field et al., 2008; Rick et al., 2019). Over the last 50 years, consumption of abalone species has shifted from wild-harvest to a 95% aquaculture-based production model, as population declines in wild fisheries have accelerated due to climate change, disease, over-harvest, starvation, and poaching (Karpov et al., 2000; Hobday et al., 2001; Neuman et al., 2010; Rogers-Bennett et al., 2010; Raemaekers, et al., 2011; Rogers-Bennett et al., 2013; De Wit et al., 2014; Cook, 2016; Lorenzo & Mantua, 2016; Rogers-Bennett & Catton, 2019). Although the majority of global aquaculture production of abalone occurs in Asia, the USA has a relatively small domestic abalone aquaculture industry based in California and Hawaii (Cook, 2016). The aquaculture of abalone and other low trophic level shellfish represents an important, expanding industry, as production requires significantly less energy input and freshwater compared to rearing terrestrial animal protein. Despite these advantages, abalone aquaculture is complex, and successful production requires the implementation of technically sophisticated and delicate culture approaches in order to successfully propagate early developmental stages that are extremely sensitive to environmental perturbation.

The acceleration of acidification along coastal regions of the west coast of North America (Gruber et al., 2012) is one feature of environmental perturbation that impacts abalone aquaculture production. For thousands of years, surface winds and a mosaic of regional currents have shaped the California Current System (CCS). The CCS seasonally transports deep-oceanic waters that are naturally low in pH into shallow nearshore environments which seasonally expose nearshore inhabitants to upwelled corrosive seawater conditions (Gruber et al., 2012; Hauri et al., 2013; Feely et al., 2016; Chan et al., 2017). This mosaic of upwelling activity within the CCS may have driven natural selection to favor OA-resilient traits in species and populations found in strong-upwelling regions. The discovery of evolved variation in resilience to OA will be beneficial for climate-smart conservation and production aquaculture programs. Climate model projections reveal outcomes with increases in strength, rate of occurrence, and duration of upwelling events, particularly along the California Coast (Gruber et al., 2012). This accelerating progression of OA within the CCS emphasizes the need to explore not only the effects of low pH during sensitive developmental stages, but also adaptation to OA and carry-over effects (Basch and Pearse 1996) from early life stages into later life history stages (Hettinger et al., 2012; Dupont et al., 2021; Lim et al., 2021; Neylan et al. 2023). Gaps in the knowledge in this area highlight the urgency to explore approaches for the diversification of aquaculture techniques to safeguard and sustain regional shellfish production. Comparing the tolerance to OA between populations residing in and out of upwelling regions will inform resource and hatchery managers’ decisions about potentially more resilient locations for broodstock collection and considerations for culture conditions onsite at abalone farms (Swezey et al. 2020).

Food availability and nutrition can mediate the negative consequences of OA on calcifying marine invertebrates (Thomsen et al., 2013). Thus, diet is likely an essential feature of aquaculture production that affects abalone sensitivity to environmental perturbations such as OA (Rossoll et al., 2012; Lemasson et al., 2019). While planktonic abalone are lecithotrophic (non-feeding), post-settlement abalone feed on the epithelial top layers of benthic crustose coralline algae (CCA) and a variety of associated microalgae species, including bacteria and diatoms. Contemporary abalone aquaculture has developed enriched diets using combinations of natural feeds (such as kelp and red algae), cultured microalgae, and artificial feeds to enhance growth and survival (Dunstan et al., 1996; Fleming et al., 1996; Dunstan et al., 2002). CCA are calcifying species of red alga known to transmit chemical cues that induce planktonic abalone to settle out of the water column and begin metamorphosis into benthic feeding snails (Morse et al., 1979; Kaspar & Mountfort, 1995; Kitting & Morse, 1997; Daume et al., 1999).

Further, CCA is known to maintain its ability to induce abalone settlement despite exposure to OA conditions (O’Leary et al., 2017). CCA tissues also provide nutrients important for successful development, such as lipids, and essential polyunsaturated fatty acids (PUFA) (Kang et al., 2013). Most abalone conservation and production aquaculture facilities in the United States use cultured diatoms, such as *Navicula* spp., which are commercially grown and readily available (Simental & Pilar, 2004); however, they do not induce the same degree of settlement success as they do not contain equivalent settlement inducing cue and generally have lower levels of essential fatty acids (EFA) and PUFA compared to CCA (Chen, 2012; Kang et al., 2013). Lipids and EFA have been shown to play an important role in development and survival of marine invertebrate larvae (Gallager et al., 1986; Kattner et al., 2003; Reitzel et al., 2004; Sewell et al., 2005), including abalone (Moran & Manahan, 2003; Swezey et al., 2020). Nevertheless, little is known about how such divergent diets may affect red abalone and their sensitivity to OA across multiple stages of development, or from different geographical sources spanning divergent, ecological and evolutionary histories.

In this study, we test for diet-OA interactions in early life stages of red abalone (*H. rufescens*). Abalone are dioecious broadcast spawners with developmental stages defined by a sensitive non-feeding, planktonic larval phase (Cox, 1962), followed by settlement to the benthos, and morphological metamorphosis (Slattery, 1992). During this transition, approximately 90% of successfully settled abalone die due to natural causes (Hahn, 1989). Studies examining the effects of OA on abalone have tended to focus on sensitivities during short windows of early development (Zippay & Hoffman, 2010; Crimm et al., 2011; Dojiri & Dojiri, 2014; Boch et al., 2017; O’Leary et al., 2017) rather than on effects that carry over long-term into later life-history stages (Hettinger et al., 2012; Hettinger et al., 2013; Neylan et al., 2023; Scanes et al., 2023). There is an urgent need to understand the capacity of red abalone to respond to OA over longer periods beyond the earliest life stages. This is because climate change is predicted to drive stronger upwelling along the California coast during the spring (Bakun et al., 2015), which will expose near-shore environments to perpetual low-pH, co-occurring with the most productive time for abalone spawning and early development (Huyer, 1983; Lynn & Simpson, 1987; Feely et al., 2008; Kelly et al., 2014; Lagos et al., 2016). It is imperative to understand the ways in which OA may disrupt abalone production, as it will be useful for informing mitigation and intervention strategies to maintain production under rapidly changing ocean conditions.

Here, we investigated how the diet of newly settled juveniles modulates the impacts of OA on nutritional quality during sensitive early developmental stages in two geographically distinct populations of red abalone; a wild-collected population from a strong upwelling region (naturally low pH) at Van Damme State Park, CA (VD), and a cultured population sourced from a zone of weak upwelling in Santa Barbara, CA (SB). We cultured red abalone from both populations from embryos to 3-months of age under either contemporary ocean pH conditions (control–pH: 8.00 pH; 450µatm CO_2_) or OA conditions (OA: 7.65 pH; 1080µatm CO_2_) that are consistent with contemporary exposures during strong upwelling events and mid-century ocean model projections for persistent conditions in red abalone benthic habitats (Gruber et al., 2012). We predicted that abalone from different populations would differ in their capacity to cope with OA, either because of physiological divergence due to natural selection in the wild or artificial selection in captivity (Hedgecock & Sly, 1990; Gaffney et al., 1996). To assess the cumulative impact of OA, we measured both ecological (survival) and physiological (total lipid quantification) parameters of larvae settled on either CCA or commercially produced *Navicula* diatoms. Compared to a *Navicula* diet, we hypothesized that the natural settlement environment and diet of CCA would reduce the energetic burden of settlement and increase overall larval energy available, allowing abalone to withstand the effects of OA. To assess the carry-over relationship between post-settlement diet and OA on juvenile abalone growth, we measured shell area of SB abalone at three months and one-year post-settlement. We discuss the implications of our results for population persistence and strategies to bolster shellfish aquaculture, given the projected increases in OA in the future.

## Methods

### Broodstock husbandry and spawning

Red abalone broodstock from The Cultured Abalone Farm (Goleta, CA) were collected in 1994 from locations throughout the Santa Barbara Channel (SB: 34.241943°N, 119.889999°W), a region that experiences weak-intermittent upwelling (Feely et al., 2016). Fourth generation descendants of this cohort were transported to University of California, Davis’ Bodega Marine Laboratory (BML) on 3 February 2016 and spawned on 5 February 2016, resulting in five maternal lineages, each crossed with pooled sperm sourced from five male abalone also from the SB population. At five to six hours post fertilization, embryos were introduced into the experimental OA system housed at BML in the early morning hours of 6 February 2016.

Between June 2015 and August 2015, 120 adult red abalone from VD, a region that experiences frequent, strong upwelling events (Feely et al., 2016), were collected by the California Department of Fish and Wildlife (CDFW) and transported to BML. These animals did not produce viable gametes during the summer in which they were collected, so they were conditioned in the lab for eleven months before spawning was again attempted. A total of four maternal lineages from multiple spawns were used in experiments. In July 2016, one male and one female produced viable gametes, which were fertilized to generate one lineage, which was introduced into the experimental OA system using the same protocol as implemented with the SB population. In February 2017, three more VD maternal lineages were generated with pooled sperm from three male VD abalone, and these animals were introduced into the experimental OA system using the same protocol.

All abalone were induced to spawn in individual buckets using the hydrogen peroxide/1M TRIS method (Tong et al., 1992; Moss et al., 1995) following an established protocol (Swezey et al., 2020). Once fertilization success was determined microscopically (percent embryos developed/ total eggs), 30,000 embryos from each maternal lineage were cultured in experimental replicates across all experiments (see experimental design and larval culturing below).

### Experimental design and larval culturing

Developing abalone were cultured in a flow-through system at 14°C and exposed to either control-pH (466 μatm *p*CO_2_; contemporary conditions of 8.00 pH, Ω_aragonite_=2) or OA (1080 μatm *p* CO_2_; near future OA 7.65, pH Ωaragonite=0.99). A complete description of the system is provided elsewhere (Swezey et al., 2017, 2020). In brief, seawater was treated in two large sumps (volume = 90 liters), which received a continuous supply of filtered UV-treated seawater and treatment air mix. Treatment air delivered to each sump was generated by blending dry CO_2_-free air with pure CO_2_ gas using digital mass flow controllers. Treatment seawater was then pumped from the sump tanks to overhead PVC manifolds, which continuously delivered seawater to the experimental units. In the first stage of the experiment, abalone were raised from fertilization until they were 7-day, planktonic veligers, in replicate larval culture chambers. Chambers were 7-L paired; circular PVC buckets fitted with 100 μm nylon mesh on the bottom to retain swimming larvae (n=30,000 per experimental unit). Treated seawater supplied to the replicate larval chambers was single pass flow-through (i.e., the source water passed through our larval chambers once and was then discarded and did not recirculate). Thus, any contamination that might have arisen in one replicate chamber (e.g., due to higher mortality of larvae in that replicate), was not circulated among the replicates within that treatment. In the second stage of the experiment, competent abalone (7 days post-fertilization [dpf]) (n=300 per experimental unit) were induced to settle using γ-aminobutyric acid (GABA) (Morse et al., 1979) and raised until 90 days post-settlement (97 dpf). These animals were housed in 120-mL experimental chambers (Starplex Scientific Leak Buster^TM^) with custom polyvinyl bottoms and fed a diet consisting of either naturally occurring CCA or a thin layer of *Navicula* diatoms (Reed Mariculture, LLC) until 97 dpf. During both stages of the experiment, seawater was sampled routinely (see below) in the sump tanks and containers to confirm that the chemical features that we manipulated were the only features of water quality that varied between treatments.

### Seawater chemistry manipulation and temperature monitoring

Seawater sampling and chemical analyses followed previously established methods (Dickinson et al., 2007). Experimental water was sampled from treatment sumps and measured three times per week throughout each experiment for pH (total scale; pH_T_) and total alkalinity (TA). These data were generated using m-Cresol purple (Shimadzu UV-1800, Shimadzu, Kyoto Japan) to spectrophotometrically quantify pH_T_. Treatment sump TA values were measured using an automated endpoint titrator (Metrohm 809 Titrando) and were standardized with certified reference material from A. Dickson at the University of California, San Diego Scripps’ Institute of Oceanography. Treatment sumps were also measured hourly for pH, salinity, and temperature using multi-parameter sondes (YSI 6920 V2, YSI, Yellow Springs, OH, USA), with pH calibrated to pH_T_ (Easley & Byrne, 2012).

To correct for minor experimental chamber variation from sump conditions, pH was sampled from three to six randomly selected units per treatment at three intervals during both the larval and post-settlement phases. Temperature offsets were calculated by averaging the temperature differences observed in experimental chambers compared to sump values as recorded by data loggers (Hobo Tidbit v2, Onset Computer, Bourne, MA, USA) that were placed in three randomly selected chambers per treatment group per diet. Differences in pH_T_ were then calculated and sump to chamber offsets were applied to the pH_T_ calibrated continuous chemistry records. For statistical comparison, calibrated hourly measurements from the continuous sump records were binned into daily experimental values. Six to ten experimental units per treatment were also randomly selected and were compared once per week to treatment sump TA values over the duration of the experiments. Because differences in TA between treatment sumps and experimental units were not detected, we concluded that negligible variation occurred; therefore, *in-situ* TA values were calculated using sump treatment values. Because TA was sampled three times per week, whereas pH_T_ was continuously measured, we regressed the established relationships of TA against salinity from continuous salinity data collected in control-pH and OA treatment sumps to establish a continuous record (Lee et al., 2000). This generated daily TA values on days where TA was not directly sampled. Regression relationships were binned and used to generate these data. These results were used to generate *p*CO_2_, DIC_calc_, Ω_aragonite_, and Ω_calcite_ values using the carbonate system software CO2SYS (Lewis & Wallace, 1998) following previously established methods (Mehrbach et al., 1973; Lee et al., 2000). For short intervals, the YSI multiparameter sondes were deployed for use in other experiments: during these times, system parameters were constructed using data from weekly sampling as opposed to continuously derived daily values and a different multi-parameter sonde (YSI Pro Plus) was used to obtain treatment sump temperature and salinity.

### Survival

Larval survival was determined at 4 dpf and 7 dpf by quantifying densities in replicate larval culture chambers (Swezey et al., 2020). To assess post-settlement survival, abalone abundance in replicate treatment units was counted under dissection microscopes at 10 dpf, 28 dpf, and 97 dpf, following Swezey et al., 2020.

### Total lipid quantification

Total lipid concentrations of red abalone larvae and newly settled abalone were determined using a modified spectrophotometric sulfophosphovanillin method (Barnes & Blackstock, 1973; Handel et al., 1985). For each larval bucket at each survival sample point, three replicate samples of 20 larvae each were collected for total lipid quantification. Swimming larvae (4 dpf & 7 dpf) and newly settled abalone (10 dpf) were randomly selected, rinsed with distilled water, pipette-dried, and stored at −80°C until analyzed following a previously established protocol (Carter et al., 2013; Swezey et al., 2020).

### Post-settlement growth analyses

Post-settlement growth rates were assessed in the SB population by photographing abalone using a stereomicroscope (Leica M125 microscope & Leica DC290 camera; Leica Microsystems, Wetzlar, Germany) at 97 dpf. After 97 dpf, SB abalone were held in ambient seawater conditions and fed *ad libitum* on a mixture of *Navicula* diatoms and dulse (*Devaleraea mollis*). Growth rates were assessed one-year post-fertilization. Images were digitally analyzed using ImageJ software v1.50i.

### Statistical analyses

We tested whether data normality (Shapiro-Wilk Test) and homogeneity of variances (Bartlett Test) met the assumptions of parametric statistical analyses (e.g. ANOVA). Statistical tests considered the three main effects of pH (control or OA), diet (CCA or *Navicula*) and source population (VD or SB), all two-way interactions, and the three-way interaction. Tukey HSD post-hoc tests were used to determine significant differences between specific treatment groups. Statistical analyses and data visualization were conducted in R Studio, version 1.1.463 (Wickam, 2016).

## Results

### Seawater chemistry and temperature

Water chemistry differed significantly between control-pH and OA treatments in each experiment (all *P*<0.0001). No variation in experimental units within these treatment groups was detected among replicate units within or between SB and VD or between exposures to CCA or *Navicula* treatments (Supplementary Table S1). Therefore, we report average treatment conditions across experiments and replicate units. During experiments, mean water temperature of the experimental units was consistent at 14.18°C across all treatments and experiments.

### Survival: Larval and post-settlement survival

Larval survival did not differ significantly between control and OA conditions within or between the SB and VD populations at both 4 dpf and 7 dpf (Supplementary Table S2; Fig. 1). In contrast, post-settlement survival at 10 dpf varied significantly between pH treatments (*P*<0.0001), settlement diets (*P*<0.0001), maternal lineages (*P*<0.0001), and for two-way interactions of pH x population (*P*=0.002), pH x maternal lineage (*P*<0.0001), settlement diet x maternal lineage (*P*<0.0001), and for the three-way interaction of pH x settlement diet x maternal lineage (*P*<0.0001) (Supplementary Table S2, Fig. 1A – D). Under control-pH conditions, SB abalone exhibited significantly greater survival when settled on CCA (mean ± SD=38.89 ± 19.46%) compared to *Navicula* diatoms (mean ± SD=22.48 ± 15.01%) (*P*<0.05) (Fig. 1A & B). This difference was also observed in the VD population under control-pH conditions, where survival was higher when settled on CCA (mean ± SD=44.34 ± 21.06%) compared to *Navicula* diatoms (mean ± SD=32.82 ± 21.45%) (*P*<0.05) (Fig. 1C & D). No significant differences in survival were observed between populations on a given settlement diet when cultured under control-pH. In contrast, when cultured under OA, survival in the SB population declined by 33% when settled on *Navicula* diatoms (compared to settlement on CCA) (*P*<0.05), but in the VD population under OA conditions no significant differences associated with settlement diet were observed.

**Fig. 1.**
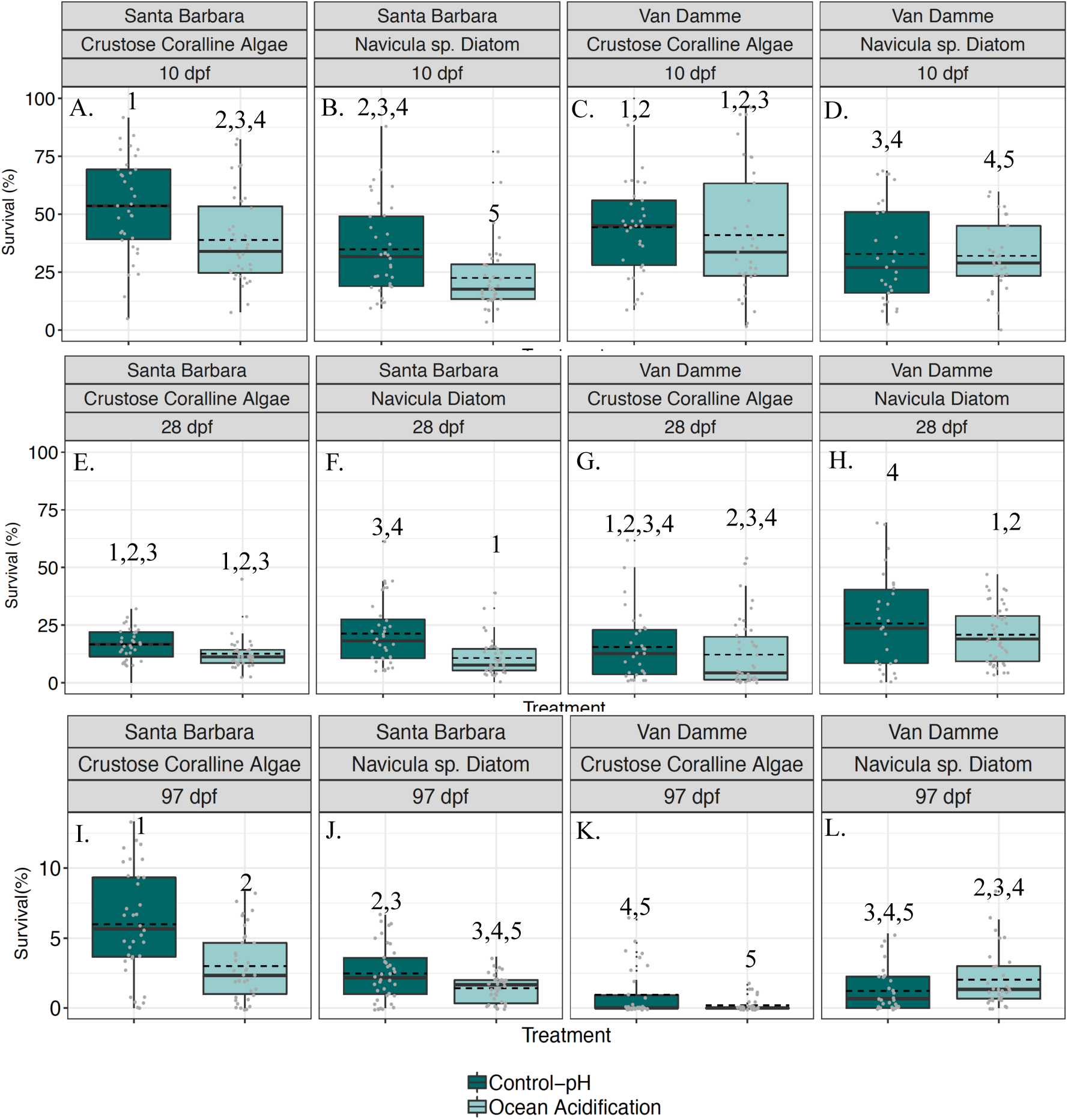
Post-settlement red abalone survival under experimental treatments. Survival under treatments was assessed at 10 days post-fertilization (dpf) (top row), 28 dpf (middle row), and 97 dpf (bottom row) for Santa Barbara (SB; first and second columns) and Van Damme (VD; third and fourth columns) populations under control-pH (dark teal) and ocean acidification (OA) conditions (light teal) when animals were raised on a diet of either crustose coralline algae (first and third columns) or *Navicula* spp. diatoms (second and fourth columns). Each post-settlement time point was statistically analyzed independently. The SB population showed elevated survival on CCA and under low CO_2_ (control-pH) conditions, whereas in the VD population, these trends diverged. Dark teal represents control-pH treatment conditions and light teal indicates ocean acidification conditions. Data points are presented as percent mean survival of replicate treatment units. Boxplot upper and lower hinges correlate with the 25^th^ and 75^th^ percentile range. Boxplot whiskers extend 1.5 times the interquartile range for each hinge. Solid black line represents the median and dashed black line represents the mean. Shared numbers within a post-settlement developmental time point indicate group means that are not significantly different based on means comparisons (*P* < 0.05).

### Survival: to one month

Red abalone survival to one month post-fertilization (28 dpf) varied significantly with pH treatment (*P*<0.0001), settlement diet (*P*<0.0031), population (*P*=0.0313), maternal lineage (*P*<0.0001), and for two-way interactions of pH treatment x settlement diet (*P*=0.0482), settlement diet x population (*P*=0.0294), pH treatment x maternal lineage (*P*<0.0002), settlement diet x maternal lineage (*P*<0.0001), and for the three-way interaction of pH treatment x settlement diet x maternal lineage (*P*=0.0082) (Supplementary Table S2; Fig. 1E – H). No significant differences were observed within each population or on a given settlement diet when cultured under control-pH conditions (Fig. 1). Within the VD population, there were no significant declines in survival on either settlement diet when cultured under OA treatment conditions (Fig 1G & 1H).

### Survival: to three months

Red abalone survival to three months post-fertilization (97 dpf) varied significantly with pH treatment (*P*<0.0001), settlement diet (*P*=0.0003), population (*P*<0.0001), maternal lineage (*P*<0.0001), and for two-way interactions pH treatment x settlement diet (*P*<0.0001), pH treatment x population (P<0.0001) settlement diet x population (*P*<0.0001), pH treatment x maternal lineage (*P*<0.0002), settlement diet x maternal lineage (*P*<0.0001), and for the three-way interaction pH treatment x settlement diet x maternal lineage (*P*=0.0225) (Supplementary Table S2; Fig.1 I – L). SB animals showed significantly greater survival when fed CCA compared to abalone fed *Navicula* when cultured under control-pH treatment and under OA conditions (*P*<0.05) (Fig. 1I & J). Survival in SB animals significantly declined when cultured under OA on each of the two diets compared to SB animals cultured under control-pH. In contrast, a decline in survival was not detected between VD abalone cultured under control-pH or OA conditions on a given diet (*P*<0.05) (Fig. 1K & L). Unlike in the SB population, under OA conditions VD abalone exhibited 148% greater survival when fed *Navicula* diatoms compared to those fed CCA (*P*<0.05). Survival did not vary among or within the populations or pH levels when comparing the response of animals from either population raised on *Navicula* at 97 dpf.

### Total lipids: Larval lipids

During the free-swimming, lecithotrophic stage, abalone larvae from VD had 29x greater total lipid concentrations compared to abalone larvae from SB (ANOVA; Supplementary Table S3, Fig. 2 A-D). During larval development the lipid concentrations of larvae from the VD population decreased by 31.3% from 4 dpf (mean ± SD=11.35 ± 3.4 ng per individual^−1^) to 7 dpf (mean ± SD=7.79 ± 5.02ng per individual^−1^) (*P*<0.05) (Fig. 2 C-D). Lipid concentrations did not vary across larval development in the SB population (Fig. 2 A-B). Larvae exposed to OA did not differ in total lipid content compared to larvae cultured under control-pH conditions in either population (Supplementary Table S3).

**Fig 2.**
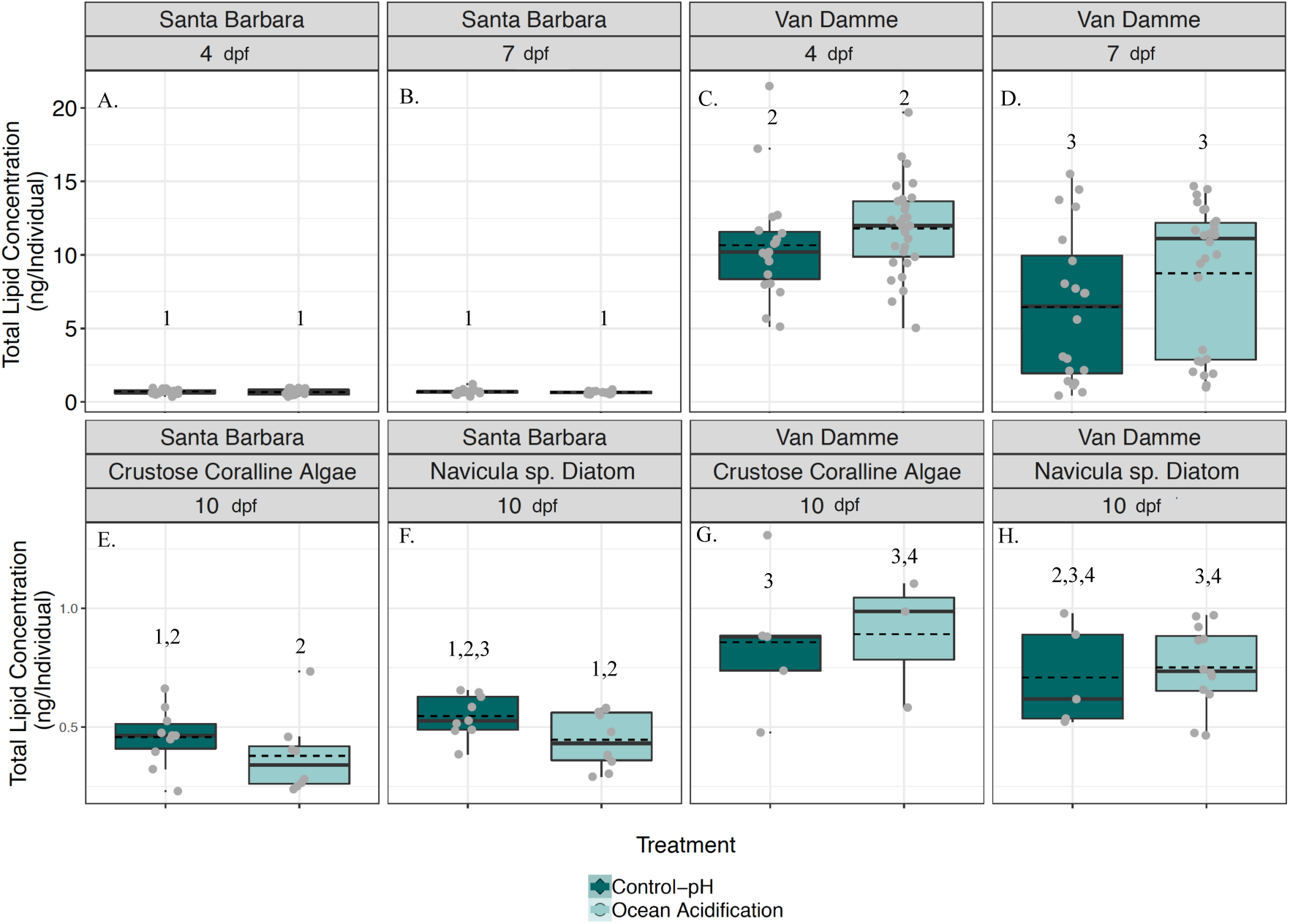
Red abalone total lipid concentrations under experimental treatments. Red abalone total lipid concentration (TLC) was determined during the free-swimming, planktonic larval phase 4 days post-fertilization (dpf) and 7 dpf (top row) as well as post-settlement at 10 dpf (bottom row) when animals were settled on diets of either crustose coralline algae (E, G) or *Navicula* spp. diatoms (F, H). Van Damme (VD) population abalone (third and fourth columns) exhibited significantly higher TLC throughout larval development compared to abalone from the Santa Barbara (SB) population (first and second columns), a trend which continued in abalone during post-settlement metamorphosis. Statistical analyses of planktonic and post-settlement TLC were assessed independently. Dark teal represents control-pH treatment conditions and light teal indicates ocean acidification conditions. Data points are presented as percent mean survival of replicate treatment units. Boxplot upper and lower hinges correlate with the 25^th^ and 75^th^ percentile range. Boxplot whiskers extend 1.5 times the interquartile range for each hinge. Solid black line represents the median and dashed black line represents the mean. Shared numbers within a post-settlement developmental time point indicate group means that are not significantly different based on means comparisons (*P* < 0.05).

### Total lipids: Post-settlement lipids

Population differences in total lipid concentrations that we observed during the larval phase carried over into the post-settlement stage (10 dpf), where VD had 69.5% greater concentrations (*P*<0.0001) of post-settlement total lipid compared to abalone from SB (VD: mean ± SD=0.78 ± 0.22 ng per individual; SB: mean ± SD=0.46 ± 0.13 ng per individual; Supplementary Table S3; Fig. 2 E – H). We observed a significant population x settlement diet interaction (*P*=0.02) such that VD abalone settled on CCA had an 89% higher lipid concentration at settlement (mean ± SD=0.87 ± 0.27 ng per individual) compared to SB abalone settled on CCA (mean ± SD=0.46 ± 0.13 ng per individual). When VD abalone were settled on *Navicula* diatoms they had 51% higher lipid concentrations (mean ± SD=0.74 ± 0.18 ng per individual) compared to SB abalone (mean ± SD=0.49 ± 0.12 ng per individual). pH treatment did not affect total lipid concentration for either population (Supplementary Table S3, Fig. 2).

### Relationship between larval lipid concentration and survival

Total larval lipid concentrations at 7 dpf were a significant predictor of survival at 10 dpf (*P*<0.0001) in the VD population and not SB population. This relationship differed depending on pH treatment (pH treatment x total larval lipids interaction; *P*<0.0001) showing a significant three-way interaction between pH, population and diet (*P*=0.0014, Supplementary Table S4; Fig. 3A – D). When settled on *Navicula* diatoms, we observed a positive relationship between survival at settlement and 7 dpf total larval lipid concentration in VD abalone when cultured under control-pH conditions. No relationship was detected when animals were cultured under OA (Fig. 3). In contrast, when abalone were raised on CCA, we observed a negative relationship between survival at settlement and 7 dpf total larval lipid concentration under OA, whereas no relationship was detected when abalone were cultured under control-pH.

**Fig. 3.**
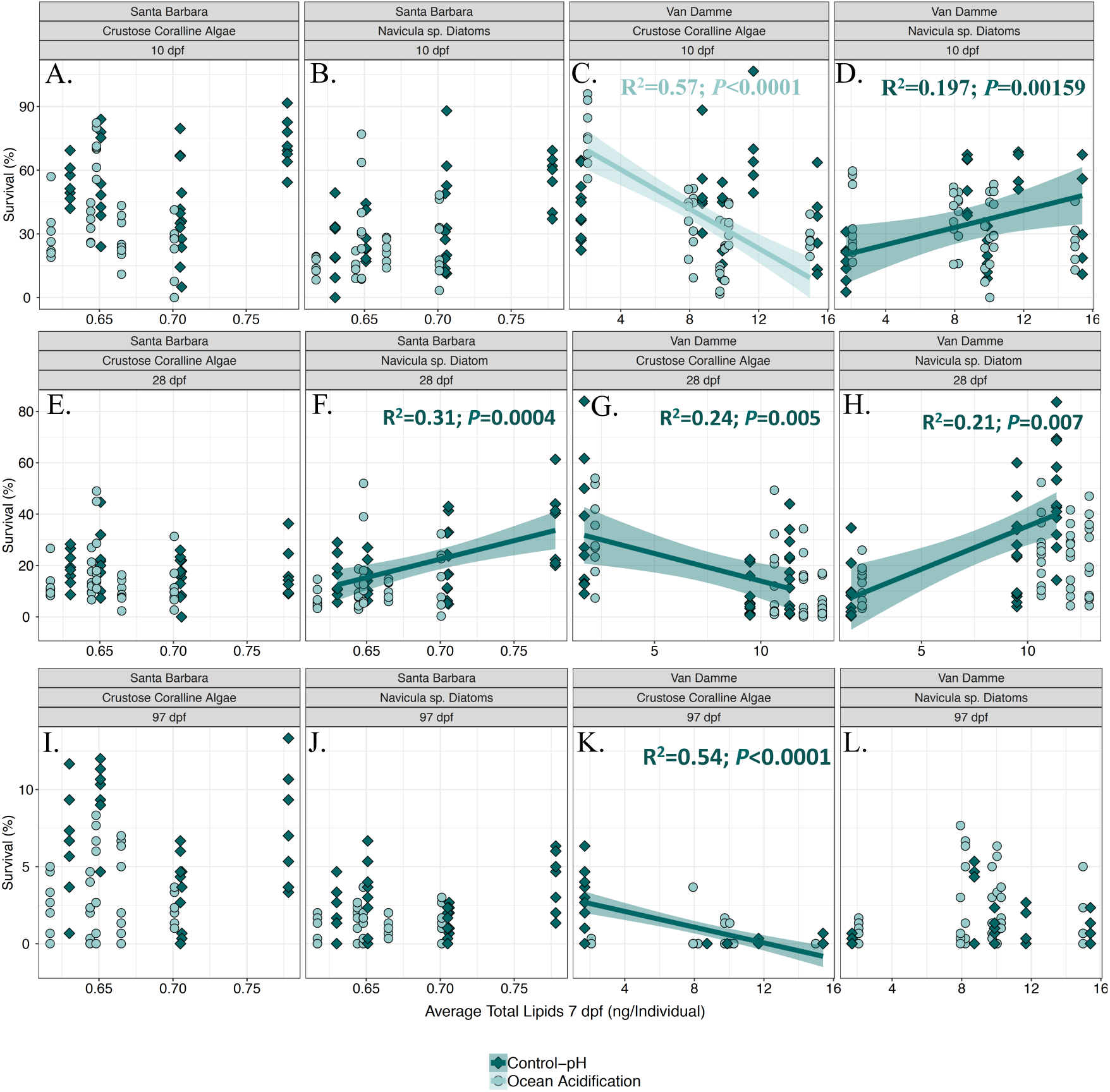
Correlation between red abalone lipid reserves and survival during post-settlement development. Lipid reserves at 7 days post-fertilization (dpf) were correlated with post-settlement survival in both Santa Barbara (SB; panels A, B, E, F, I, J) and Van Damme (VD; panels C, D, G, H, K, L) populations, although this effect was much more pronounced in the VD population. Regression lines show significant relationships between lipids and survival under control-pH (dark teal diamonds) or ocean acidification conditions (light teal circles) at 10, 28, and 97 days post-fertilization (dpf; top, middle, and bottom rows, respectively) on diets of coralline algae (first and third columns) and *Navicula* spp. diatoms (second and fourth columns). Shaded areas around regression lines represent the 95% confidence band. Summary statistics associated with regressions are provided in Table 5.

Total larval lipid concentrations at 7 dpf were also a significant predictor of survival at 28 dpf (*P*=0.00002), though this correlation varied depending on settlement diet (settlement diet x total larval lipid interaction; *P*=0.00002), pH treatment (pH treatment x total larval lipid interaction; *P*=0.03), and varied depending on population (population x total larval lipid interaction; *P*=0.04), including a significant three-way interaction (pH x population x diet *P*=0.00003; Supplementary Table S4; Fig. 3E – H). At 28 dpf, 7 dpf lipid concentrations had no relationship to survival in SB animals raised on CCA, whereas VD animals exhibited a strong negative relationship (control-pH: *P*=0.005; OA: *P*<0.0001; (Fig. 3). SB animals displayed a significant positive relationship between total larval lipid concentrations at 7 dpf and survival at 28 dpf under both control-pH and OA conditions when fed *Navicula* diatoms (control-pH: *P*=0.0004; OA: *P*=0.023). In contrast, VD abalone fed *Navicula* diatoms exhibited a significant positive relationship under control pH (*P*=0.007), but no relationship under OA was detected (Fig. 3). A significant negative relationship was observed between 7 dpf lipids and 28 dpf survival in the VD population when animals were raised on CCA under OA conditions.

Total larval lipid concentrations at 7 dpf were a significant predictor of survival at 97 dpf (*P*=0.00001); however, this observed relationship differed depending on pH treatment (pH treatment x total larval lipid interaction; *P*=0.00001), by settlement diet (settlement diet x total larval lipid interaction; *P*=0.02), including a significant three-way interaction (*P*=0.07) (Fig. 3 I-L). Survival at 97 dpf tended to show a strong (VD control pH, P<0.0001), to weak negative relationship with 7 dpf lipid when raised on CCA versus weak positive relationships to survival in both populations when raised on *Navicula* diatoms (Fig. 3).

### Cumulative diet effects on survival

When survival and total larval lipid concentration data points at 7 dpf from both populations were combined, larval lipid concentrations were a significant predictor of survival at all time points (all *P*<0.0001; Fig. 4). This relationship varied by settlement diet (settlement diet x total larval lipid interaction; *P*<0.0001, for all developmental stages), and by pH treatment at 97 dpf (pH treatment x total larval lipid interaction; *P*<0.0001) (Supplementary Table S5; Fig. 4). We observed a consistent negative relationship between 7 dpf lipid concentrations and survival across time points when animals were settled on CCA, versus a consistently positive relationship between survival and lipid concentration when animals were raised on *Navicula* diatoms (Fig. 4 A-F) except for at 97 dpf, where survival under control-pH showed no significant relationship to lipid concentration at 7 dpf in animals raised on *Navicula* (Fig. 4).

**Fig 4.**
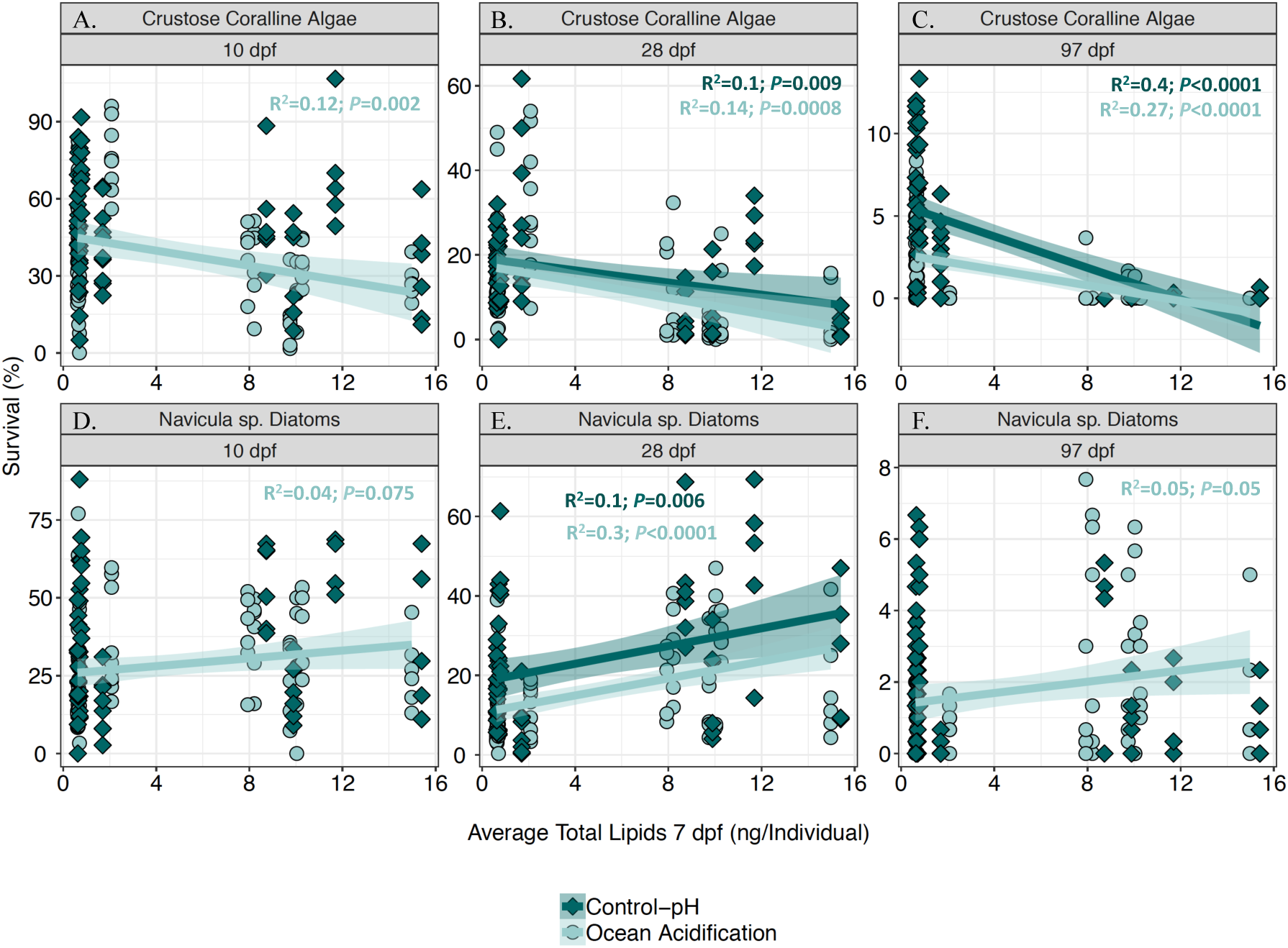
Correlation between red abalone post-settlement survival and total lipid concentration (TLC) for both populations combined under experimental conditions. Animals were raised on either crustose coralline algae (top row) or *Navicula* spp. diatoms (bottom row) at 10 days post-fertilization (dpf), 28 dpf, and 97 dpf (first, second, and third columns, respectively). Dark teal represents control-pH conditions and light teal indicates ocean acidification conditions. When raised on coralline algae, red abalone show a negative relationship between survival and TLC, whereas this relationship becomes positive when abalone are raised on diatoms. Shaded areas around the slopes of the lines represent 95% confidence bands. Only statistically significant predictors are highlighted in each respective panel (P<0.05). Summary statistics associated with regressions are provided in Table 8.

### Carry-over diet effects on Santa Barbara juvenile shell growth three months after settlement

Juvenile shell growth was measured only in SB animals. Shell growth of abalone at three months post-settlement varied significantly with pH treatment (*P*<0.0001) and by settlement diet (*P*<0.0001) (Supplementary Table S6; Fig. 5). For red abalone settled on CCA, average shell areas for animals cultured under OA conditions (mean ± SD=3.6 mm ± 1.9 mm^2^) were 20% larger than those cultured under control-pH conditions (mean ± SD=2.99 mm^2^ ± 1.6 mm^2^) (*P*<0.05) (Fig. 5). For red abalone cultured under control-pH, those settled on *Navicula* diatoms were 36.6% smaller (mean ± SD=1.5 mm^2^ ±1.6 mm^2^) compared to abalone settled on CCA (*P*<0.05) (Fig. 5).

**Fig 5.**
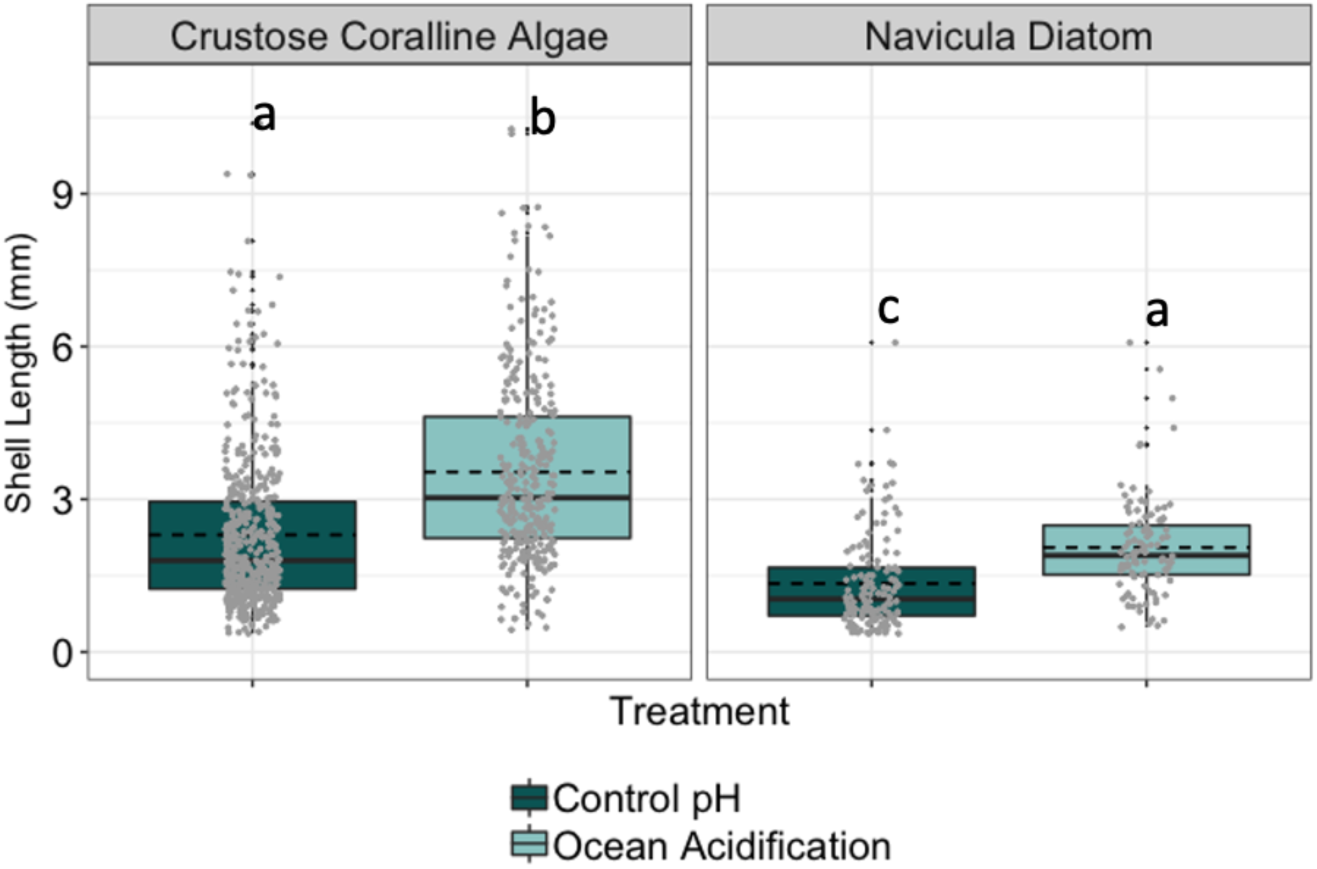
Shell lengths of red abalone sourced from the Santa Barbara (SB) population were determined three months post-settlement (97 dpf) when settled on either crustose coralline algae (left panel) or *Navicula* spp. diatoms (right panel). Dark teal represents control-pH conditions and light teal indicates ocean acidification conditions. Data points are presented as individual abalone shell measurements for each treatment group. Boxplot upper and lower hinges correlate with the 25^th^ and 75^th^ percentile range. Boxplot whiskers extend 1.5 times the interquartile range for each hinge. Solid black line represents the median and dashed black line represents the mean. Shared letters indicate group means that are not significantly different based on means comparisons (*P*<0.05).

### Carry-over diet effects on Santa Barbara juvenile shell growth one year after settlement

Settlement diet significantly affected shell growth, where abalone settled on a diet of CCA had significantly larger shells (mean ± SD=170.3 ± 78.49 mm^2^) compared to abalone settled on *Navicula* diatoms (mean ± SD=116.05 ± 71.34 mm^2^) one-year post-settlement under ambient conditions (*P*<0.001; Table 6, Fig. 6). No differences in shell size were detected between control-pH and OA conditions within the settlement diet at one year of age under ambient ocean conditions (Fig. 6).

**Fig 6.**
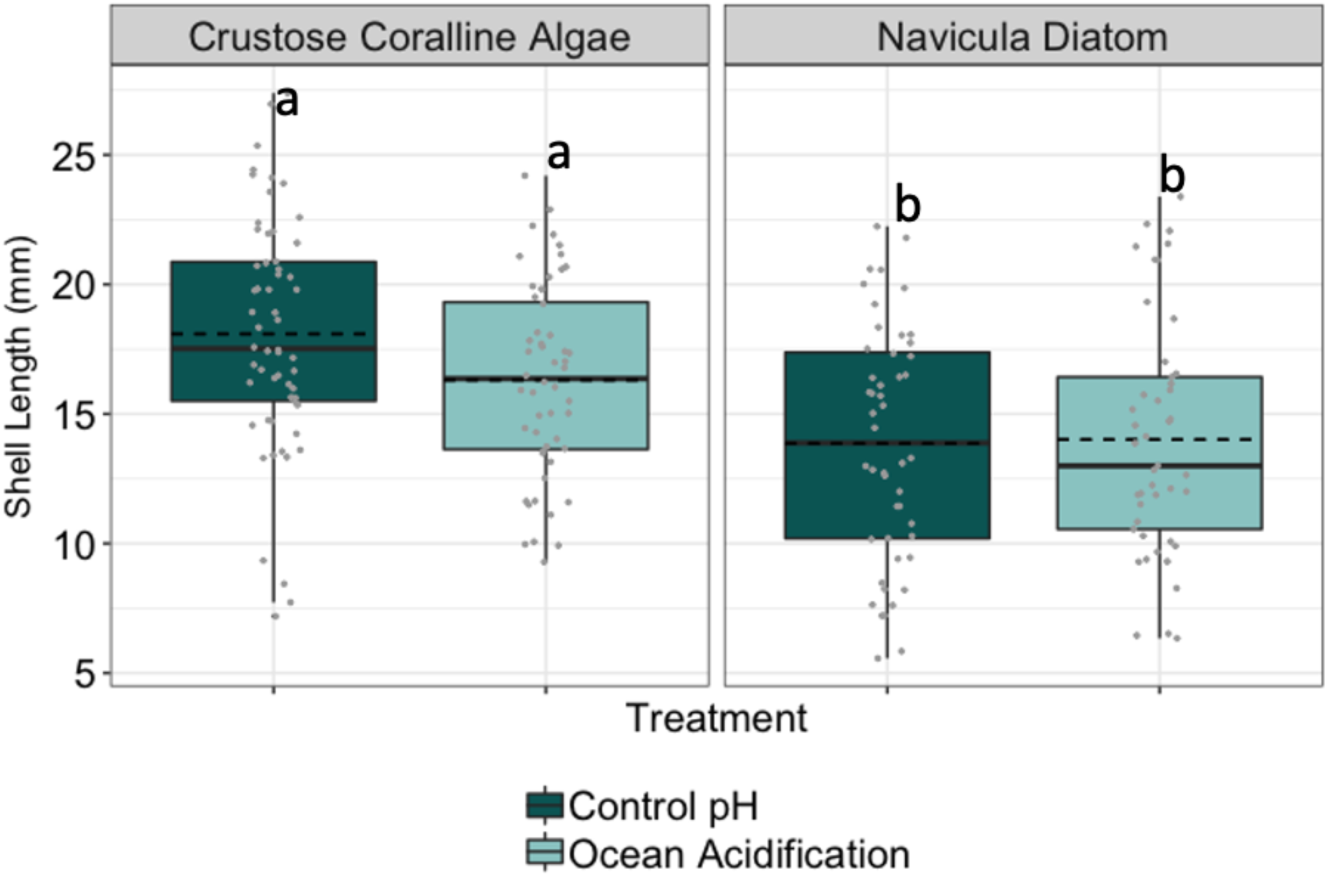
Red abalone growth at 12 months. Red abalone shell lengths from the Santa Barbara (SB) population were measured at 12 months post-fertilization in animals originally settled on either crustose coralline algae (left panel) or *Navicula* sp. diatoms (right panel). Abalone settled and raised on CCA achieved larger sizes by one year of age compared to animals raised on diatoms. Dark teal represents control-pH treatment conditions and light teal represents ocean acidification conditions. Data points are presented as individual abalone shell measurements for each treatment group. Boxplot upper and lower hinges correlate with the 25^th^ and 75^th^ percentile range. Boxplot whiskers extend 1.5 times the interquartile range for each hinge. Solid black line represents the median and dashed black line represents the mean. Shared letters indicate group means that are not significantly different based on means comparisons (*P* < 0.05).

## Discussion

In order to inform strategies for the management of wild and aquaculture raised populations of abalone, we investigated whether diet and source population modify the response of the red abalone to OA in early life history phases. We also tested whether the source population, including both a wild population from a strong upwelling region (VD) and a captive farm population (SB) from a weak-intermittent upwelling region, would modify these associations between diet and the impacts of OA on production. Finally, we examined the carry-over effect of settlement diet on juvenile growth in the captive farm population (SB). Notable results from this study included: (1) CCA initially enhanced survival during the earliest stages of development in both populations under both control-pH and OA conditions, though this effect remained significant in only the SB population by 97 dpf; (2) SB animals showed a marked decline in survival under OA across post-settlement time points, whereas survival in VD animals was not affected by OA; (3) diet modified the relationship between post-settlement survival and total larval lipid concentrations; (4) the relationship between larval lipid concentrations and post-settlement survival was more pronounced for the VD population than the SB population although strong contrasting trends between diet types were observed when data from both populations was combined (5) SB juvenile shell growth was enhanced by OA treatment exposure; however, (6) after one year, only the effects of settlement diet impacted SB juvenile shell growth.

### Diet, pH treatment, and population affect post-settlement survival

Our results highlight the complexity of factors affecting red abalone post-settlement survival including source population, diet, and OA. Larval settlement on CCA initially enhanced post-settlement survival in both populations compared to abalone settled on *Navicula*. Differences in early diet may affect the nutritional state of sensitive post-settlement abalone and ultimately their survival. In addition to providing settlement cues and substrate for larval abalone (Morse et al., 1979; Shepard & Turner, 1985; Kitting & Morse, 1997; Daume et al., 1999; Suenaga et al., 2004; Kang et al., 2014), CCA offers enriched polyunsaturated fatty acids (PUFA) compared to *Navicula* (Chen, 2012; Kang et al., 2013). The digestibility and assimilation of essential fatty acids (EFA) in *Haliotis* spp. varies by the particular class of these molecules, with PUFA being the most digestible EFA compared to saturated fatty acids (SFA) and monounsaturated fatty acids (MUFA) (Dunstan et al., 2002). Greater dietary availability of these molecules can markedly impact sensitive life history stages, such as during post-settlement and metamorphosis, which are known to be energetically demanding stages for marine invertebrates (Bogan et al., 2019). We posit that differences in nutritional quality associated with differences in the fatty acid profiles of these diets may contribute to the OA-buffering effect of CCA, which is especially magnified in SB animals that started life with very low available lipid energy reserves. Findings from Rossell et al. (2012) suggests that OA can also reduce the ratio of PUFA:SFA in diatoms, with resulting negative effects on invertebrate consumers. This may further help explain the particularly acute negative effects of OA on the lipid-poor SB population when raised on *Navicula*.

In both populations, we found that survival of post-settlement abalone exposed to OA was enhanced by CCA at 10 dpf compared to survival on *Navicula* diatoms (Fig. 1 A-D). These results suggest that CCA provides a more favorable settlement environment for abalone compared to *Navicula* diatoms, which afforded some protection from OA-induced impacts on survival. The initial OA-buffering effect of CCA on survival observed in the SB population persisted to a lesser degree at 28 dpf and emerged again by 97 dpf (Fig. 1 I-L). We know that the settlement cue of CCA is resilient to OA (O’Leary et al., 2017). However, this effect was not detected in the VD population at the later post-settlement stages, indicating that CCA did not evoke the same beneficial outcome under OA in VD abalone.

Population differences observed for post-settlement survival in response to diet may be attributed to differences in nutritional requirements between t wild and captive populations of red abalone. Differences in biochemical composition and dietary needs have been shown to exist between wild and cultured populations (Gross et al., 1998; Karapanagiotidis et al., 2006), including abalone (Dunstan et al., 1996). Newly settled abalone in the wild feed on a wide variety of biofilms, CCA, filamentous algae, bacteria and diatoms. In contrast, cultured abalone feed on a narrow range of diatoms, less diverse bacteria and artificial feeds under typical farm conditions. Therefore, adaptation to captive conditions may have contributed to evolved nutritional differences and enhanced the OA-buffering effect of CCA on the SB abalone compared to the VD population. Farmed red abalone may experience nutritional deficiencies during sensitive early stages because controlled access to settlement substrates on farms provides a diet that is not typical of those available in natural habitats. As such, CCA may mitigate these deficiencies in cultured abalone, especially in the context of energetic stress linked to OA.

The difference in the OA-buffering effect of CCA between VD and SB abalone may also be a result of local adaptation between strong and weak upwelling habitats; these habitat differences provide different multi-generational histories of exposure to low pH during early development. Previous research has demonstrated populations of marine species can be adapted to local environmental conditions (Thor & Oliva, 2015; Calosi et al., 2017; Gaitan-Espitia et al., 2017), such as coastal upwelling zones (Padilla-Gamiño et al., 2016; Evans et al., 2017; Swezey et al., 2017; Griffiths et al., 2019). Adaptation to strong seasonal upwelling conditions on the Mendocino coast may have also contributed to evolved differences in OA-buffering capacity previously described between the two geographically distinct populations (Swezey et al., 2020). More population contrasts (multiple cultured and wild populations, from multiple strong and weak upwelling regions) are necessary to more definitively identify the causes of the population differences observed and how they drive sensitivity to the dietary buffering of OA impacts.

### Relationship between total larval lipids and post-settlement survival

The settlement and survival enhancing properties of CCA have been well documented in the literature for a variety of species (Morse et al., 1979; Kitting & Morse, 1997; Daume et al., 1999; Laimek et al., 2008; Gómez-Lemos et al., 2018). In this study, we provide additional nuance by demonstrating that CCA and *Navicula* diets differentially modulate the complex relationship between larval lipid concentrations and post-settlement survival. Abalone with greater total larval lipid concentrations at 7 dpf had higher survival when settled and fed on *Navicula* diatoms, whereas animals with lower total larval lipid concentrations at the completion of the larval phase had greater survival when settled and fed CCA. This relationship between larval lipid concentrations and survival was also modified by OA and varied between populations (Fig. 3 & 4), which suggest that a complex interaction between genetics and environmental conditions govern physiology and early development in red abalone.

We found that VD abalone had much greater total lipid content compared to SB abalone prior to settlement, though this difference diminished by 10 dpf (Fig. 2). This suggests that these populations differ in how they store and mobilize lipid reserves during development. Previous work has shown that wild and cultured Australian green-lip abalone (*H. laevigata*) differ in tissue lipid compositions as adults (relative concentrations of MUFA, PUFA, and SFA) and that this variation is linked to differential experimental diets (Dunstan et al., 1996). Here, we show that a similar plasticity may exist in abalone at the earliest development stages. While the exact mechanism driving this variation is unclear, this variation has consequences for survival under OA (Dunstan et al., 1996). It remains to be determined whether the population differences in lipid profiles observed in the present study are a result of different evolutionary histories between strong and weak upwelling habitats, or because of more recent divergence in captivity compared with wild stocks or other mechanisms of plasticity.

Considering data from both populations together, we detected a consistent positive relationship between larval lipids and post-settlement survival when animals were settled and raised on *Navicula* diatoms under OA conditions (Fig. 4); however, this trend was not significant for abalone cultured under control-pH at 10 dpf and 97 dpf indicating that OA stress strongly structures this predictive relationship. When abalone were fed a natural diet of CCA, a consistent negative relationship between larval lipids and post-settlement survival was observed across pH treatments (Fig. 4) suggesting that the abalone are not nutrient limited when settled on CCA.

When examining the relationship between total larval lipids at 7 dpf and post-settlement survival for each population separately, we observed additional complexity. Elevated larval lipid concentrations were associated with survival on *Navicula* in the VD population under control-pH, whereas lower starting lipid concentrations were associated with greater survival on CCA under OA conditions (Fig. 3). In the SB population, these differences were absent, possibly because these abalone started life with such low lipid concentrations, or other nutritional requirements that differed as a result of aquaculture-based selection of the broodstock (Nettleton & Exler, 1992; Dunstan et al., 1996).

The survival outcomes observed in the present study may also be due to the negative effects of OA on lipid metabolism, which could limit energy allocation for responding to persistent stress (Lee et al., 2018). Environmental stress affects energetic demand, diverting energy required for growth and development to compensate for increased metabolic needs (Sokolova et al., 2013). The two populations may experience (energetically) the same water temperature very differently. Under ambient pH conditions, the Palourde clam (*Ruditapes decussatus*) displayed a negative relationship between lipid content and survival similar to the response observed in red abalone settled on CCA (Matias et al., 2011), indicating the animals were not nutrient limited. However, this relationship was modified by feeding developing Palourde clams a mixed diet of CCA and *Navicula* diatoms (Matias et al., 2011). Our observed positive relationship between lipid concentration and survival on *Navicula* under OA may be attributed to elevated energy demands required for coping with pH stress when raised on this diet (Leung et al., 2020), which may be especially pronounced as OA alters the nutritional quality of diatoms (Rossoll et al., 2012). Endogenous lipid reserves are critical for molluscan larval development and survival (Moran & Manahan, 2003; Hettinger et al., 2013), which may help to explain why abalone with greater lipid stores had a greater proportion of survivors compared to abalone with fewer lipid reserves. These results suggest that, compared to CCA, diatoms may provide insufficient nutrition to maintain homeostasis when cultured under OA conditions.

### Diet and pH affect SB juvenile growth

Many studies show that the nutritional quality of diets available to developing marine invertebrate larvae can mitigate the negative effects of environmental stress on growth (Hettinger et al., 2013; Asnaghi et al., 2014; Bogan et al., 2019). For example, juvenile purple sea urchins (*Paracentrotis lividus*) fed CCA had enhanced growth rates compared to purple urchins fed non-calcifying macroalgae under OA conditions (Asnaghi et al., 2014), similar to the growth patterns observed in the present study (Fig. 5 & 6). Here, the effect of early maturation on CCA also enhanced juvenile shell growth in SB cultured red abalone; however, this growth-enhancement effect of CCA varied with pH, and the effect of early life pH treatment and was no longer discernable at one-year post-settlement (Fig. 6). Impacts to shell growth affect time to reach size refuge from predation in the wild and market size in aquaculture production. In the present study, we confirm that diet manipulation can ameliorate the negative effects of OA on both yield and growth, consistent with findings from other studies (Hettinger et al., 2013; Asnaghi et al., 2014).

This study contributes to a growing body of research highlighting the importance of diet nutritional quality during early development in wild and cultured molluscs. Alternative diets directly affect growth, settlement, metamorphosis, and the ability to respond to stress (Waldock & Nascimento, 1979; Enright et al., 1986; Delaunay et al., 1993; Soudant et al., 1998; Aranda-Burgos et al., 2014).

Exploration of the complex and divergent nutritional and metabolic demands of wild and cultured shellfish during sensitive developmental stages will increase the ability of managers and aquaculture practitioners to better understand and mitigate for the negative consequences of OA. Future work should explore the effect of OA on the fatty acid profiles of red abalone across multiple life history stages, as there is a growing awareness that population variation (due to random-neutral drift, or natural selection in the wild or captivity) may affect how diet and OA interact to impact development, physiology, growth, survival, and production yields. This work will ultimately prove essential to informing effective mitigation/intervention strategies needed to enhance nutritional state and promote survival in early life stages of red abalone and other shellfish under changing ocean chemistry. Efforts to maintain sustainable populations in both commercial and conservation production aquaculture efforts are critical given the precipitous global decline of abalone (Peters et al., 2024). Further exploration of the complex and interacting dietary and energetic requirements of commercially produced marine invertebrate larvae under ocean climate change will help advance both the conservation and production aquaculture of valuable genus *Haliotis* spp. worldwide.

## Compliance With Ethical Standards

The authors have no relevant financial or non-financial interests to disclose.

## Acknowledgements

We sincerely thank L. Heidenreich, G. Ayad, S. Bashevkin, A. Broffman, S. Garcia, M. Heard, E. Hubbard, T. Leung, S. Merolla, K. Magaña, P. Shukla, N. Sippl-Swezey, C. Vines, B. Walker, H. Weinberger, and T. Winquist for their assistance in experimental research and spawning. We are grateful for the encouragement and advice from C. Catton, B. Gaylord, and T. Hill. We thank A. Todgham for the use of research equipment. We also thank the California Department of Fish and Wildlife for the collection of wild abalone from Van Damme State Park. This research was funded by California Sea Grant R/HCME-10, the California Conservation Genomics Project, and by the National Oceanic and Atmospheric Administration Small Business Innovation Research Contracts WC-133R-15-CN-0072 and WC-133R-16-CN-0105 to The Cultured Abalone Farm.

## Data Availability

The datasets generated during and/or analyzed during the current study are available in a supplementary appendix (Supplementary Appendix S1).

---

**Supplementary Table S1.** Summary of larval phase and post-settlement seawater carbonate chemistry. Means, standard deviations and the statistical significance of differences between treatments (columns 2-4) for carbonate system parameters are presented. Total alkalinity (TA), pH total (pH_T_), salinity, and temperature were used to calculate the partial pressure of CO_2_ (*p*CO_2calc_), aragonite saturation state (Ωaragonite), and total dissolved inorganic carbon (DIC). Tukey-Kramer tests were used to determined significant differences. Shared letters in the “Significant Grouping” column indicate group means (for the treatments represented in the prior four columns) that are not significantly different based on means comparisons (*P*<0.05).

| Seawater Parameter<br>Larval Phase | SB Control-pH | SB - OA | VD Control-pH | VD - OA | Significance<br>Grouping |
| --- | --- | --- | --- | --- | --- |
| Temperature (°C) | 14.679±0.05 | 14.51±0.09 | 14.413±0.26 | 14.532±0.28 | A,A,A,A |
| Salinity | 32.483±0.17 | 32.203±0.36 | 31.786±1.48 | 32.277±1.42 | A,A,A,A |
| TA (μmol Kg<sub>w</sub><sup>-1</sup>) | 2208±8.45 | 2207±9.19 | 2181±55.7 | 2203±37.7 | A,A,A,A |
| DIC (μmol Kg<sub>w</sub><sup>-1</sup>) | 2032±13.4 | 2160±12.7 | 1972±59.6 | 2150±45 | A,B,A,B |
| pH<sub>T</sub> | 7.997±0.03 | 7.636±0.03 | 8.091±0.05 | 7.653±0.06 | A,B,C,B |
| $p\text{CO}_{2\text{calc}}$ | 445±33.4 | 1116±85.3 | 346±46 | 1075±166 | A,B,A,B, |
| $\Omega_{\text{aragonite}}$ | 2.02±0.11 | 0.949±0.06 | 2.34±0.17 | 0.987±0.12 | A,B,C,B |
| $\Omega_{\text{calcite}}$ | 3.16±0.11 | 1.49±0.06 | 3.67±0.17 | 1.55±0.12 | A,B,C,B |
| Seawater Parameter<br>Post-settlement Phase |  |  |  |  |  |
| Temperature (°C) | 14.230±0.314 | 14.074±0.442 | 14.202±0.433 | 14.216±0.359 | A,B,A,B |
| Salinity | 32.752±0.557 | 32.833±0.592 | 33.191±1.071 | 33.544±0.796 | A,A,B,C |
| TA (μmol Kg<sub>w</sub><sup>-1</sup>) | 2215.4±0.557 | 2217.4±19.9 | 2213±63.5 | 2215±70.6 | A,A,A,A |
| DIC (μmol Kg<sub>w</sub><sup>-1</sup>) | 2056±23.4 | 2168±26.4 | 2032±63.9 | 2158±77.2 | A,B,C,B |
| pH<sub>T</sub> | 7.960±0.0387 | 7.640±0.0503 | 8.005±0.069 | 7.659±0.061 | A,B,C,B |
| $p\text{CO}_{2\text{calc}}$ | 492±48.3 | 1107±125 | 440±82.5 | 1057±155.0 | A,B,C,B |
| $\Omega_{\text{aragonite}}$ | 1.86±0.174 | 0.960±0.121 | 2.06±0.296 | 1.02±0.162 | A,B,C,B |
| $\Omega_{\text{calcite}}$ | 2.92±0.269 | 1.50±0.189 | 3.22±0.463 | 1.59±0.252 | A,B,C,B |

**Supplementary Table S2.** Statistical (ANOVA) analyses of experimental treatment effects on survival across developmental time points. Treatment effects on survival were assessed at 10 dpf, 28 dpf and 97 dpf (three consecutive tables below) in response to three main effects including population (Santa Barbara or Van Damme), pH treatment (control-pH or ocean acidification), and settlement diet (crustose coralline algae or *Navicula* sp. diatoms). Models tested for interactions between main effects, and maternal lineage within population was also modeled.

| ANOVA | d.f. | SS | *F* | *P* |
| --- | --- | --- | --- | --- |
| **Effects on Post-Settlement Survival (%) to 10 dpf** |  |  |  |  |
| pH Treatment | 1 | 0.401 | 18.12 | <0.0001 |
| Settlement Diet | 1 | 1.19 | 53.72 | <0.0001 |
| Population | 1 | 0.009 | 0.4 | 0.5 |
| pH Treatment x Population | 1 | 0.22 | 9.76 | 0.002 |
| Settlement Diet x Population | 1 | 0.03 | 1.37 | 0.24 |
| pH Treatment x Settlement Diet x Population | 1 | 6.7e-6 | 0.0003 | 0.98 |
| pH Treatment x Maternal Lineage [Population] | 6 | 2.47 | 18.63 | <0.0001 |
| Settlement Diet x Maternal Lineage [Population] | 6 | 1.22 | 9.17 | <0.0001 |
| pH Treatment x Settlement Diet x Maternal Lineage [Population] | 6 | 0.5 | 3.75 | 0.0014 |
| Maternal Lineage [Population] | 6 | 1.7 | 12.96 | <0.0001 |
| **Effects on Post-Settlement Survival (%) to 28 dpf** |  |  |  |  |
| ANOVA | d.f. | SS | *F* | *P* |
| pH Treatment | 1 | 0.2 | 18.17 | <0.0001 |
| Settlement Diet | 1 | 0.09 | 8.95 | 0.0031 |
| pH Treatment x Settlement Diet | 1 | 0.04 | 3.9 | 0.048 |
| Population | 1 | 0.05 | 4.69 | 0.03 |
| Settlement Diet x Population | 1 | 0.05 | 4.8 | 0.029 |
| pH Treatment x Settlement Diet X Population | 1 | 0.01 | 1.23 | 0.27 |
| pH Treatment x Maternal Lineage [Population] | 6 | 0.3 | 4.67 | 0.0002 |
| Settlement Diet x Maternal Lineage [Population] | 6 | 1.18 | 17.98 | <0.0001 |
| pH Treatment x Settlement Diet x Maternal Lineage [Population] | 6 | 0.19 | 2.96 | 0.0082 |
| Maternal Lineage [Population] | 6 | 0.31 | 4.85 | 0.0001 |
| **Effects on Post-Settlement Survival (%) to 97 dpf** |  |  |  |  |
| ANOVA | d.f. | SS | *F* | *P* |
| pH Treatment | 1 | 636.5 | 22.66 | <0.0001 |
| Settlement Diet | 1 | 374.7 | 13.34 | 0.0003 |
| pH Treatment x Settlement Diet | 1 | 418.2 | 14.89 | 0.0001 |
| Population | 1 | 2603.5 | 92.68 | <0.0001 |
| pH Treatment x Population | 1 | 476.2 | 16.95 | <0.0001 |
| Settlement Diet X Population | 1 | 1636.9 | 58.27 | <0.0001 |
| pH Treatment x Population x Settlement Diet | 1 | 3.45 | 0.12 | 0.72 |
| Maternal Lineage [Population] | 6 | 1643.5 | 9.75 | <0.0001 |
| pH Treatment x Maternal Lineage [Population] | 6 | 760.1 | 4.51 | 0.0002 |
| Settlement Diet x Maternal Lineage [Population] | 6 | 1036.7 | 6.15 | <0.0001 |
| pH Treatment x Settlement Diet x Maternal Lineage [Population] | 6 | 423.06 | 2.51 | 0.02 |

**Supplementary Table S3.** Summary statistics (ANOVA) for experimental treatment effects on total lipid concentration across three developmental time points, including 4 dpf, 7 dpf, and 10 dpf (three tables below). Main treatment effects included settlement diet (crustose coralline algae or *Navicula* sp. diatoms – for 10 dpf treatment only), pH treatment (control-pH or ocean acidification), and population (Santa Barbara or Van Damme). Models tested for interactions between main effects, and maternal lineage within population was also included in the model.

| **Effects on Total Lipid Concentration at 4 dpf** |  |  |  |  |
| --- | --- | --- | --- | --- |
| ANOVA | d.f. | SS | *F* | *P* |
| pH Treatment | 1 | 6.6 | 0.06 | 0.8 |
| Population | 1 | 5663.2 | 54.35 | <0.0001 |
| pH Treatment x Population | 1 | 6.03 | 0.06 | 0.8 |
| Maternal Lineage [Population] | 6 | 22.9 | 0.35 | 0.9 |
| pH Treatment x Maternal Lineage [Population] | 6 | 422.7 | 0.67 | 0.7 |
| **Effects on Total Lipid Concentration at 7 dpf** |  |  |  |  |
| ANOVA | d.f. | SS | *F* | *P* |
| pH Treatment | 1 | 0.02 | 0.003 | 0.9 |
| Population | 1 | 933.7 | 159.9 | <0.0001 |
| pH Treatment x Population | 1 | 0.002 | 0.0002 | 0.9 |
| Maternal Lineage [Population] | 6 | 874.6 | 24.9 | <0.0001 |
| pH Treatment x Maternal Lineage [Population] | 6 | 38.9 | 1.11 | 0.4 |
| **Effects of Total Lipids Concentration at 10 dpf** |  |  |  |  |
| ANOVA | d.f. | SS | *F* | *P* |
| pH Treatment | 1 | <0.01 | 0.14 | 0.71 |
| Population | 1 | 1.58 | 55.54 | <0.0001 |
| Settlement Diet | 1 | <0.01 | 0.01 | 0.91 |
| pH Treatment x Population | 1 | 0.02 | 0.79 | 0.38 |
| pH Treatment x Settlement Diet | 1 | <0.01 | 0.02 | 0.9 |
| Population x Settlement Diet | 1 | 0.16 | 5.53 | 0.02 |
| pH Treatment x Population x Settlement Diet | 1 | <0.01 | 0.03 | 0.87 |

**Supplementary Table S4.** Significance of predictive relationships between larval lipids and post-settlement survival under experimental conditions. R^2^ values and the significance of regressions between larval lipids concentrations and post-settlement survival at 10 dpf, 28 dpf, and 97 dpf for SB and VD populations cultured on crustose coralline algae (CCA) or diatoms (NV) under either control-pH or OA.

| Regression Analysis | $R^2$ | $F (df1,df2)$ | $P$ |
| --- | --- | --- | --- |
| **10 dpf Survival (Fig. 3A-D)** |  |  |  |
| SB-CCA-Control | 0.06 | 2.18 (1,34) | 0.15 |
| SB-NV-Control | 0.25 | 11.07 (1,33) | 0.002 |
| SB-CCA-OA | 0.04 | 1.18 (1,33) | 0.28 |
| SB-NV-OA | 0.05 | 1.77 (1,36) | 0.19 |
| VD-CCA-Control | 0.00 | 0.01 (1,31) | 0.94 |
| VD-NV-Control | 0.2 | 6.61 (1,27) | 0.02 |
| VD-CCA-OA | 0.57 | 56.51 (1,42) | <0.0001 |
| VD-NV-OA | 0.02 | 1.00 (1,44) | 0.32 |
| **28 dpf Survival (Fig. 3E-H)** |  |  |  |
| Regression Analysis | $R^2$ | $F (df1,df2)$ | $P$ |
| SB-CCA-Control | 0.08 | 2.70 (1,33) | 0.11 |
| SB-NV-Control | 0.31 | 15.36 (1,35) | 0.0004 |
| SB-CCA-OA | 0.02 | 0.74 (1,35) | 0.39 |
| SB-NV-OA | 0.14 | 5.61 (1,35) | 0.02 |
| VD-CCA-Control | 0.24 | 8.98 (1,29) | 0.005 |
| VD-NV-Control | 0.21 | 8.41 (1,31) | 0.007 |
| VD-CCA-OA | 0.41 | 5.61 (1,39) | <0.0001 |
| VD-NV-OA | 0.06 | 2.76 (1,42) | 0.1 |
| **97 dpf Survival (Fig. 5I-L)** |  |  |  |
| SB-CCA-Control | 0.02 | 2.70 (1,33) | 0.41 |
| SB-*NV*-Control | 0.06 | 2.27 (1,36) | 0.14 |
| SB-CCA-OA | 0.001 | 0.04 (1,33) | 0.8 |
| SB-*NV*-OA | 0.08 | 3.08 (1,35) | 0.08 |
| VD-CCA-Control | 0.54 | 38.8 (1,33) | <0.0001 |
| VD-*NV*-Control | 0.04 | 1.21 (1,26) | 0.28 |
| VD-CCA-OA | 0.00 | 0.02 (1,41) | 0.9 |
| VD-*NV*-OA | 0.012 | 0.74 (1,40) | 0.4 |

**Supplementary Table S5.** Effects summaries of cumulative red abalone post-settlement survival under experimental treatments and the relationship between total lipid concentration and survival under pH (control vs. OA) and diet treatments (CCA, NV) of abalone combined from VD and SB populations. R^2^ and p-values represent the strength of regression relationships between larval lipid concentrations and post-settlement survival at 10 dpf, 28 dpf, and 97 dpf.

| ANOVA |  | d.f. | SS | F | P |
| --- | --- | --- | --- | --- | --- |
| Effects on 10 dpf Survival |  |  |  |  |  |
| pH Treatment |  | 1 | 0.48 | 12.29 | 0.0005 |
| Settlement Diet |  | 1 | 1.12 | 29.05 | <0.0001 |
| pH Treatment X Settlement Diet |  | 1 | 0.02 | 0.43 | 0.51 |
| Total Lipids |  | 1 | 0.03 | 0.86 | 0.35 |
| pH Treatment X Total Lipids |  | 1 | 0.05 | 1.29 | 0.25 |
| Settlement Diet X Total Lipids |  | 1 | 0.59 | 15.39 | 0.0001 |
| pH Treatment X Settlement Diet X Total Lipids |  | 1 | 0.03 | 0.75 | 0.39 |
| Effects on 28 dpf Survival |  |  |  |  |  |
| pH Treatment |  | 1 | 0.21 | 13.64 | 0.0003 |
| Settlement Diet |  | 1 | 0.18 | 11.87 | 0.0007 |
| pH Treatment X Settlement Diet |  | 1 | 0.05 | 3.14 | 0.07 |
| Total Lipids |  | 1 | 0.01 | 0.73 | 0.39 |
| pH Treatment X Total Lipids |  | 1 | 0.003 | 0.24 | 0.62 |
| Settlement Diet X Total Lipids |  | 1 | 0.67 | 34.11 | <0.0001 |
| pH Treatment X Settlement Diet X Total Lipids |  | 1 | 0.03 | 0.11 | 0.73 |
| Effects on 97 dpf Survival |  |  |  |  |  |
| pH Treatment |  | 1 | 0.007 | 15.72 | <0.0001 |
| Settlement Diet |  | 1 | 0.003 | 7.33 | 0.0072 |
| pH Treatment X Settlement Diet |  | 1 | 0.005 | 11.52 | 0.0008 |
| Total Lipids |  | 1 | 0.02 | 48.58 | <0.0001 |
| pH Treatment X Total Lipids |  | 1 | 0.007 | 15.85 | 0.0001 |
| Regression Analysis |  | R<sup>2</sup> | F (df1,df2) |  | P |
| 10 dpf Survival Fig. 6A & D |  |  |  |  |  |
| CCA-Control |  | 0.03 | 1.72 (1,67) |  | 0.2 |
| NV-Control |  | 0.03 | 2.2 (1,62) |  | 0.14 |
| CCA-OA |  | 0.12 | 10.34 (1,77) |  | 0.002 |
| NV-OA |  | 0.04 | 3.23 (1,35) |  | 0.07 |

| **28 dpf Survival Fig. 6B & E** |  |  |  |
| --- | --- | --- | --- |
| CCA-Control | 0.1 | 7.27 (1,64) | 0.009 |
| *NV*-Control | 0.1 | 8.02 (1,65) | 0.006 |
| CCA-OA | 0.14 | 12.18 (1,76) | 0.0008 |
| *NV*-OA | 0.3 | 20.77 (1,79) | <0.0001 |
| **97 dpf Survival Fig. 6C & F** |  |  |  |
| CCA-Control | 0.4 | 47.75 (1,68) | <0.0001 |
| *NV*-Control | 0.04 | 2.33 (1,64) | 0.13 |
| CCA-OA | 0.27 | 29.45 (1,78) | <0.0001 |
| *NV*-OA | 0.05 | 3.95 (1,77) | 0.05 |

**Supplementary Table S6.**
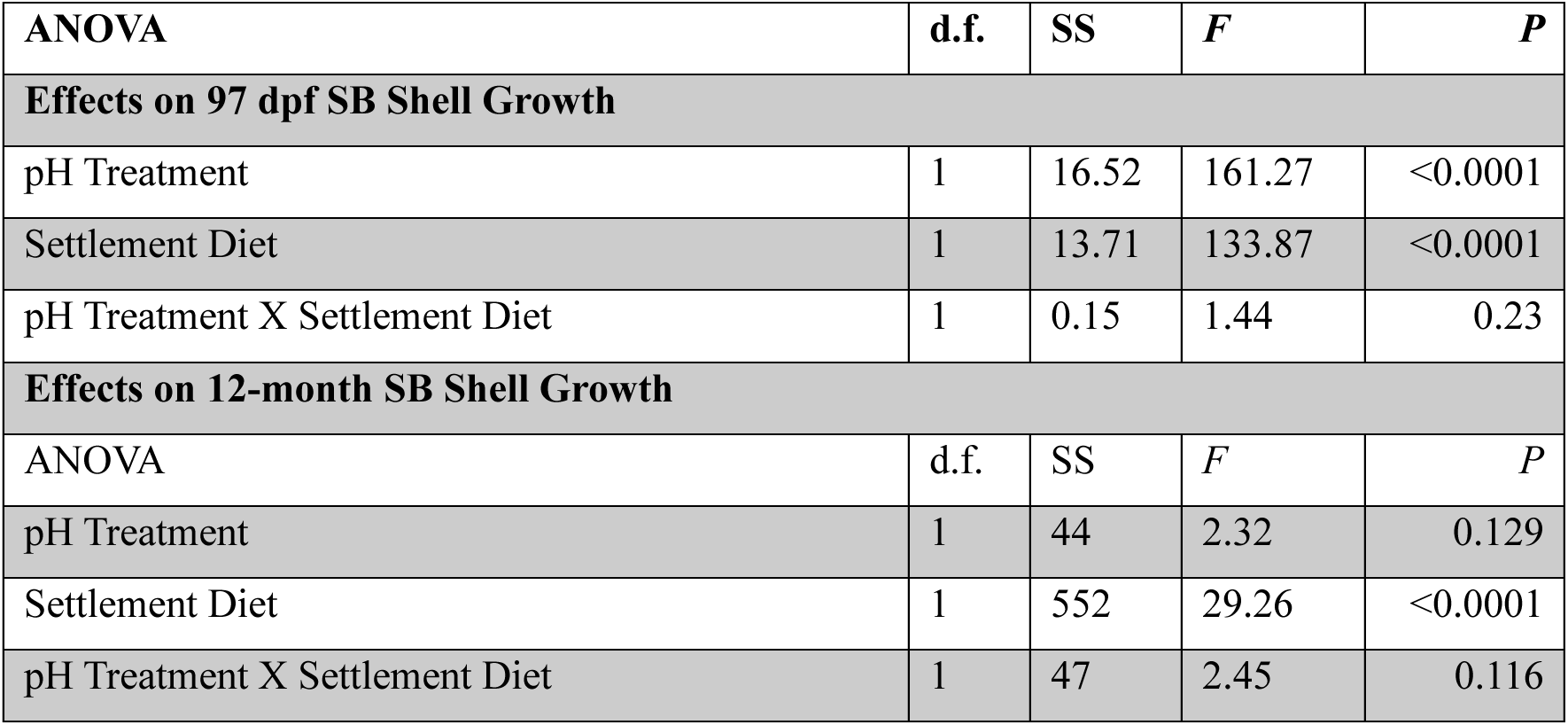
Summary of treatment effects on post-settlement shell growth at 97 days post-fertilization (dpf) and 12 months post-fertilization. Settlement diet and pH treatment modulate post-settlement shell growth for abalone settled on either crustose coralline or diatoms differentially across assessment time points.

| ANOVA | d.f. | SS | *F* | *P* |
| --- | --- | --- | --- | --- |
| **Effects on 97 dpf SB Shell Growth** |  |  |  |  |
| pH Treatment | 1 | 16.52 | 161.27 | <0.0001 |
| Settlement Diet | 1 | 13.71 | 133.87 | <0.0001 |
| pH Treatment X Settlement Diet | 1 | 0.15 | 1.44 | 0.23 |
| **Effects on 12-month SB Shell Growth** |  |  |  |  |
| ANOVA | d.f. | SS | *F* | *P* |
| pH Treatment | 1 | 44 | 2.32 | 0.129 |
| Settlement Diet | 1 | 552 | 29.26 | <0.0001 |
| pH Treatment X Settlement Diet | 1 | 47 | 2.45 | 0.116 |

